# Maternal malnutrition impacts placental morphology and transport. An origin for poor offspring growth and vulnerability to disease

**DOI:** 10.1101/727404

**Authors:** Kristin L Connor, Mark Kibschull, Elzbieta Matysiak-Zablocki, Tina Tu-Thu Ngoc Nguyen, Stephen G Matthews, Stephen J Lye, Enrrico Bloise

**Affiliations:** Lunenfeld-Tanenbaum Research Institute, Mount Sinai Hospital, Toronto, Ontario, Canada; Health Sciences, Carleton University, Ottawa, Ontario, Canada; Department of Physiology, University of Toronto, Toronto, Ontario, Canada; Department of Obstetrics and Gynaecology, University of Toronto, Toronto, Ontario, Canada; Department of Morphology, Federal University of Minas Gerais, Belo Horizonte, Brazil

**Author notes:** Correspondence: Dr. Kristin Connor, Department of Health Sciences, Carleton University, 1125 Colonel By Drive, 3310 Health Sciences Building, Ottawa, ON, Canada.

**Keywords:** placenta, development, malnutrition, transport, morphology

## Abstract

The placenta promotes fetal growth through nutrient transfer and selective barrier systems. An optimally developed placenta can adapt to changes in the pregnancy environment, buffering the fetus from adverse exposures. We hypothesised that the placenta adapts differently to suboptimal maternal diets, evidenced by changes in placental morphology, developmental markers, and key transport systems. Mice were fed a control diet (CON) during pregnancy, or undernourished (UN) by 30% of control intake from gestational day (GD)5.5-18.5, or fed 60% high fat diet (HF) eight weeks before and during pregnancy. At GD18.5, placental morphometry, development, and transport were assessed. Junctional and labyrinthine areas of UN and HF placentae were smaller than CON by >10%. Fetal blood space area and fetal blood space:fetal weight ratios were reduced in HF vs. CON and UN. Trophoblast giant cell marker *Ctsq* mRNA expression was lower in UN vs. HF, and expression of glycogen cell markers *Cx31.1* and *Pcdh12* was lower in HF vs. UN. Efflux transporter *Abcb1a* mRNA expression was lower in HF vs. UN, and *Abcg2* expression was lower in UN vs. HF. mRNA expression of fatty acid binding protein *Fabp*_*pm*_ was higher in UN vs. CON and HF. mRNA and protein levels of the lipid transporter FAT/CD36 were lower in UN, and FATP4 protein levels were lower in HF vs. UN. UN placentae appear less mature with aberrant transport. HF placentae adapt to excessive nutrient supply. Understanding placental adaptations to common nutritional adversities may reveal mechanisms underlying the developmental origins of later disease.

## Introduction

The placenta is an active participant in pregnancy, acting as a conduit between the mother and fetus. In this capacity, it can promote fetal growth through nutrient and oxygen transfer^1–3^, and minimise adverse exposures from reaching the fetus via efflux transporters and other barrier systems^4^. When its function is optimal, the placenta can adapt to changes in the pregnancy environment and buffer the fetus from adverse exposures^1,5^. But if the placenta’s ability to respond to environmental stimuli is compromised, perhaps due to altered placental development and/or function, adaptation may be insufficient or absent, and fetal growth and development may be adversely affected^1,5^. Poor fetal growth is problematic: it is associated with suboptimal growth trajectories in early life and increased risk for chronic diseases long-term^6,7^. Yet, it remains unclear how the placenta adapts to altered pregnancy environments in a manner that could shape fetal growth and possibly underlie early origins of later disease.

In order for the placenta to function optimally, it first needs to develop appropriately. Key aspects of placental development include differentiation of cell types and placental structures^8^, which are involved initially in the establishment and maintenance of pregnancy, and later in nutrient and gas exchange between mother and fetus. Placental morphology is exquisitely regulated by molecular events, with some of the best studied placental differentiation and development markers identified in the trophoblast lineage in the mouse placenta^8–10^. In the junctional zone (JZ), the major endocrine layer of the mouse placenta, the trophoblast giant cells (TGCs) (in part identified by expression of the gene *Prl3b1*) are differentiated cells that maintain contact with the maternal environment throughout development^11^ and are critical for many maternal adaptations during pregnancy. This includes regulation of blood flow and synthesis of placental lactogen hormones^12^. Spongiotrophoblast cells (identified by expression of *Tpbpa*), situated between TGCs and the inner labyrinth layer, make up the JZ of the mouse placenta, providing structural support for the growth of the labyrinthine villi and limit fetal endothelium overgrowth^13^. Within the spongiotrophoblast layer, glycogen trophoblast cells (in part identified by expression of *Cx31.1* and *Pcdh12*) develop and migrate into the decidua from mid gestation and assemble around spiral arteries to provide endocrine and/or structural support to exchange structures^9,14,15^. Substantial expansion of these glycogen cells^16^ between E14.5-18.5 results in a significant increase in glucose storage for the fetus, likely contributing to fetal growth in late gestation and providing a source of energy for the fetus during delivery^14,16,17^. In the labyrinth zone, the major murine placental exchange layer, there exists a layer of sinusoidal giant cells (with some level of exchange function^11^, identified by *Prl3b1* and *Ctsq* gene expression), and two layers of syncytiotrophoblast that are connected by gap junctions^18^. When genes regulating placental differentiation and development are absent or non-functional^10,12,19^, placental development is impeded, and fetal development can be altered^20–22^.

Secondary to structure, the selective permeability of the placenta is critical to fetal growth and development^1^. The fetus is reliant on transport of nutrients, particularly glucose, amino acids, and fatty acids, via the placenta through nutrient transporters and blood flow regulation^2,23^. Fatty acids are particularly important for fetal brain development, cellular signalling^24^, and adipose tissue expansion, and their transfer to the fetus increases with advancing gestation, such that peak adipose deposition occurs at term^25^. Fatty acids are shuttled across the placenta typically down a maternal-fetal concentration gradient via membrane fatty acid transport proteins (FATPs), CD36, and fatty acid binding protein plasma membrane (FABP-pm) ^26–28^. Additionally, throughout pregnancy, placental efflux transporters (namely ATB-binding cassette transporters, or ABC transporters) are critical for removing waste and limit the accumulation of maternally-derived exogenous substrates in the syncytiotrophoblast barrier, thereby serving to protect the fetus against factors in the maternal circulation^4,29^. They do this by transferring endogenous substrates from the cytosol to the extracellular space, or from one side of a barrier to another^30^. Placental P-glycoprotein (P-gp, encoded by the *ABCB1* gene) and breast cancer related protein (BCRP, *ABCG2*) are two major ABC transporters in the placenta and show opposing expression profiles across human gestation: *ABCB1* mRNA and P-gp protein levels are high in first trimester, decreasing towards term^31^, whilst BCRP protein levels increase towards term, with mRNA levels not changing across gestation^32,33^. Some ABC transporters are critical for lipid transport, and many are sensitive to the effects of lipids, which appear to influence ABC transporter activity^34^, although these relationships are poorly defined in pregnancy. These selective barrier functions of the placenta have been described in isolation in normal pregnancy^33,35–38^ and in response to various adverse exposures including malnutrition^39–41^ and maternal infection or (meta)inflammation^29,42^. However, the impact of adverse exposures on the function of nutrient supply and xenobiotic efflux has not been assessed concomitantly. It is important to comprehensively understand how the placenta functions as a conduit to regulate fetal development, since there are many situations wherein adverse exposures like malnutrition, infection/inflammation and/or medication use coexist^43–47^.

Not all fetuses exposed to adverse environments *in utero* show signs of compromised growth, which may be explained by the placenta’s adaptive capacity, influencing its ability to act as a selective barrier for trophic factors and other xenobiotics. We hypothesised that the placenta could adapt differently to maternal UN and HF diets, through changes in its development and function, and these adaptations would explain why offspring of UN and HF mothers grow differently *in utero*, and could reveal mechanisms underpinning developmental origins of later disease. To address this, we assessed select placental developmental markers, and ABC and fatty acid transporters at the end of pregnancy in dams fed a normal diet, UN, or fed a HF diet, and determined the effects of these diets on maternal metabolism and fetal growth.

## Methods

### Animal model

All experiments were approved by the Animal Care Committee at Mount Sinai Hospital. In brief, male and female C57BL/6 mice were obtained from Jackson Laboratories and housed in a single room under constant temperature of 25°C with a 12:12 light-dark cycle and free access to food and water. Females were randomised to one of three nutritional groups^48,49^: i) mice fed a control diet (Dustless Precision Pellets S0173, BioServe, Frenchtown, NJ, USA) ad libitum before mating and throughout pregnancy (CON, n=7); ii) or mice fed a control diet ad libitum before mating and until gestational day (GD) 5.5, and then undernourished (UN) restricting food intake by 30% of control intakes^50^ from GD 5.5-17.5, after which females were fed control ad libitum for the remainder of the study (n=7); iii) or mice fed a high fat (HF) diet (60% kcal as fat, D12492, Research Diets, New Brunswick, NJ, USA) ad libitum from 8 weeks before mating and throughout pregnancy (n=8). All breeding males were fed control diet ad libitum for the duration of the study. Mating occurred at ∼10 weeks of age, at which time pregnant females were housed individually in cages with free access to water and their respective diets. This day represented day 0.5 of pregnancy (GD0.5; term = 19 days). All dams were weighed and food intakes recorded weekly before pregnancy and daily during pregnancy.

### Biospecimen collection and processing

At GD18.5, the period of peak fetal growth and just following attainment of peak placental volume, maternal blood space, and fetal capillary development^51^, dams were killed by cervical dislocation. Immediately following, tail glucose was measured using a commercial glucose meter (Roche Accucheck). Dams were decapitated and trunk blood was collected into heparin-coated tubes to isolate plasma. Fetuses and placentae were rapidly dissected from the uterus and weighed, and tissues were flash frozen in liquid nitrogen and stored at −80C, for later molecular analyses, or rinsed with ice-cold 1X PBS, then fixed in 10% neutral-buffered formalin, followed by storing in 70% EtOH at 4°C before being embedded in paraffin for later histological analyses.

### Histological analyses

5 µm paraffin embedded placental sections were stained according to standard protocols with haematoxylin and eosin Y (H&E), Periodic acid–Schiff (PAS), or with specific primary and secondary antibodies as described below. H&E and PAS-stained slides were digitally scanned at 5x or 20x magnification, respectively, using a Hamamatsu digital slide scanner at the Lunenfeld-Tanenbaum Research Institute Optical Imaging Facility. The resultant images were viewed using ImageJ and the ImageJ NDPITools plugins^52^. Measures of placental junctional and labyrinth zone areas were calculated as described by Bloise et al.^53^ (2012). PAS-positive cells were counted in the entire placental section and expressed as the number of PAS-positive cells/spongiotrophoblast area or /JZ area. Placental area and image analysis scoring was conducted on one male and one female placentae from each dietary group where each sample selected represented the average fetal and placental weight for that dietary group^53,54^. One observer blinded to the experimental groups performed the image analysis.

Sections from each placentae were also stained for Ki67 P-gp and BCRP. For Ki67 staining, sections were first rehydrated and then quenched using 0.3% hydrogen peroxide (Fisher, Toronto, ON, Canada) in methanol for 30 minutes at room temperature. Sections underwent antigen retrieval first by boiling in 1X unmasking solution (DAKO, Mississauga, ON, Canada) for 30 minutes and next by boiling in 10 mM sodium citrate solution for 8 minutes. Sections were blocked using serum-free protein blocking solution (DAKO) for one hour at room temperature and incubated overnight at 4°C with 1:200 dilution of mouse anti-Ki67 antibody (Novocastra, Concord, ON, Canada) in antibody diluent (DAKO). Next, a 1:100 dilution of goat biotinylated anti-mouse antibody (Vector Labs, Burlington, ON, Canada) in antibody diluent (DAKO) was applied for 1 hour at room temperature, followed by a 1:2000 dilution of streptavidin-horseradish peroxidase (Invitrogen, Burlington, Canada) in 1X PBS for 1 hour at room temperature. DAB peroxidase substrate (Vector Labs, Burlingame, CA, USA) was applied for 1 minute 30 seconds to visualise the antibody signals. Between all steps sections were washed 3 times with PBS with 0.1% Tween20. Sections were counterstained with Gill’s #1 haematoxylin. Monoclonal mouse IgG1 antibody (DAKO) served as the negative control. Six images were randomly captured at 40X magnification (Leica DMIL LED inverted microscope and QCapture Pro software) in each of the following three placental regions for each placenta: chorionic plate, spongiotrophoblast, labyrinth zone. Image analysis was conducted by a single observer blinded to the experimental groups by counting the number of immunoreactive Ki67-positive cells per image.

Placentae were stained for P-gp (1:500, D-11 Santa Cruz, Mississauga, Canada) and BCRP (1:200, Calbiochem, Etobicoke, ON, Canada) as described above with the following changes: sections were quenched using 0.03% hydrogen peroxide in 1X PBS for 30 minutes at room temperature; secondary antibody was goat biotinylated anti-mouse antibody (1:200, Vector Labs); and DAB peroxidase substrate incubation time to visualise antibody signals was 30 seconds for both P-gp and BCRP. From stained sections, six images were randomly captured at 20X magnification (Leica DMIL LED inverted microscope and QCapture Pro software) in the spongiotrophoblast and in the labyrinth zones. Semi-quantitative analysis to score staining intensity in each image was undertaken by a single observer blinded to the experimental groups as described previously^55^. Staining was scored as absent (0), weak (1), moderate (2), strong (3) and very strong (4). Mean staining intensity across the six images for each placental zone for each placenta was calculated.

### in situ hybridisation

Placental architecture at GD18.5 was analysed using the Visiopharm NewCAST software program (Horsholm, Denmark) following in situ hybridisation using the following probes: *Ctsq, Prl3b1, Pcdh12*, and *Tpbpa* as described previously^19^. 20% of the area per placental section was randomly counted at 20x magnification (using an Olympus BX61 microscope) with 20 points per visual field, representing 8.9 mm^2^ per point/count. A total of 3 or 4 sections per placenta were analysed from 4 female and 4 male placentae per dietary group. Total or relative areas for specific structures were calculated for each section and averaged per placenta. Structures of interest included glycogen cells, PAS-positive cells, sinusoidal TGCs, TGCs, interstitial glycogen cells, maternal blood space, fetal blood space, maternal blood space in the JZ, labyrinthine trophoblast and spongiotrophoblast area. JZ size was calculated by summing spongiotrophoblast, TGC and JZ sinus areas. LZ size was calculated by summing fetal blood space, labyrinthine sinus and labyrinthine trophoblast areas.

### Plasma biomarker measurements

Commercially available mouse-specific plate assays were used according to manufacturer’s instructions to measure biomarkers in maternal circulation. Plasma insulin (Ultrasensitive Mouse ELISA, ALPCO, Salem, NH, USA), leptin (Mouse ELISA, Crystal Chem, Downers Grove, IL, USA), adiponectin (Mouse High Molecular Weight ELISA, ALPCO, Salem, NH, USA), and triglycerides (LabAssay Triglyceride Kit, Wako, Richmond, VA, USA) were measured in plasma. Free fatty acids (Free Fatty Acid Quantitation Kit, Sigma-Aldrich, St. Louis, MO, USA) were measured in maternal erythrocytes pooled for each dietary group due to the small volume of erythrocytes available from each animal making individual animal measures impossible.

### mRNA isolation and expression in placenta

Total RNA was extracted from placenta^29^ using TRIZOL reagent (Invitrogen) following manufacturer’s instructions. RNA purity and concentration were assessed by spectrophotometric analysis (Nanodrop) and RNA integrity was verified using gel electrophoresis. 1 µg RNA was reverse transcribed using 5X iScript Reverse Transcription Supermix (BioRad, Mississauga, ON, Canada). A non-reverse transcription (NRT; absence of enzyme) sample was reverse transcribed to provide a negative control for the RT reaction in downstream PCR applications.

Real-time quantitative (q) PCR (Bio-Rad CFX384, Hercules, CA, USA) was used to measure relative expression of the following ABC and fatty acid transport genes: *Abcb1a* and *Abcb1b* (encoding P-gp), *Abcg2* (encoding BCRP), *Fatp1, Fatp4* (encoding fatty acid transport proteins [FATP], part of the solute carrier family 27 [SLC27A]), *Fat/Cd36* (fatty acid translocase), *Fabppm* (encoding fatty acid binding protein plasma membrane), the placental lipases^56, 57^ *El* (encoding endothelial lipase) and *Lpl* (encoding lipoprotein lipase), and for placental developmental markers: *Ctsq, Prl3b1, Pcdh12, Tpbpa, Cx31.1*. Primer sequences for genes of interest and three stably expressed reference genes (*β-actin, Hprt-1, Gapdh* for ABC and fatty acid genes; *Tbp, Hprt-1, Gapdh* for developmental genes) are listed in Supplementary Table 1). A non-template control (NTC; absence of cDNA template) was prepared as a negative control for the PCR reaction. Each real-time qPCR reaction mix was prepared with SYBR Green JumpStart Taq ReadyMix (Sigma, Oakville, ON, Canada) and forward and reverse primer mix (3 µM) for each gene. Standard curves for each gene were run. Standards, samples and controls were run in triplicate under the following cycling conditions: 95°C for 20s; 40 cycles of 95°C for 5s and 60°C for 20s and a melt curve at 65°C→95°C with a 0.5°C increment every 5s. Relative gene expression for each sample was calculated using the Cq value of the gene of interest relative to the geometric mean of the reference genes’ Cq values using Pfaffl’s relative ratio^58^.

### Protein isolation and expression in placenta

Total protein was extracted from placenta using RIPA buffer with cOmplete™ protease inhibitor (Sigma, Oakville, ON, Canada) and 100mM sodium orthovanadate. Protein concentration was determined using BCA assay (Pierce, Thermo Fisher, Mississauga, ON, Canada).

For western blot experiments, 75 µg of each protein was separated by electrophoresis (100V, 1 hour) using 9% SDS polyacrylamide gels and then transferred to a 0.2µM polyvinylidene fluoride (PVDF) membrane over 10 minutes (Bio-Rad Trans-Blot Turbo). Membranes were blocked with 5% skim milk in 0.05% TBS-T for 1 hour. Membranes were then incubated with primary antibodies overnight at 4°C. The primary antibodies used were: rabbit anti-FATP1 (1:250, abcam); rabbit anti-FATP4 (1:1000, abcam), rabbit anti-FAT/CD36 (1:500, abcam,) rabbit anti-FABPpm (1:1000, abcam); goat anti-β-actin (1:500 used to normalise FATP1, FATP4 and FAT/CD36, or 1:1000 used to normalise FABPpm, Santa Cruz). PVDF membranes were subsequently incubated for 1 hour at room temperature with donkey HRP-linked anti-rabbit secondary antibody (1:5000 for FATP1, FATP4, FAT/CD36; 1:10000 for FABPpm, GE Healthcare, Bio-Science, Baie d’Urfe, QC, Canada) or donkey anti-goat secondary antibody (1:5000 for β-actin, Santa Cruz). Protein-antibody complexes were detected following incubation of the membrane with SuperSignal West Femto (Thermo Fisher). Chemiluminescence was detected under UV (VersaDoc, Bio-Rad). Intensity of the bands of interest were quantified using Image Lab software (Bio-Rad), where the percent intensity for each band (for proteins of interest and β-actin) was calculated from the volume intensity of each band, and percent intensity of the bands from the proteins of interest signals were normalised against those from the β-actin signal.

For placental LPL protein levels, a mouse-specific ELISA was used (Cusabio, Houston, TX, USA) according to manufacturer’s instruction. All protein samples were normalised to the same concentration in sample diluent prior to assay. Samples were run in duplicate and mean (SD) %CV between duplicates was 6.98 ± 4.58%.

### Placental cytokine measurements

Protein isolated from placental homogenates were assayed for cytokine levels according to manufacturer’s instructions using the Bio-Plex Pro Mouse Cytokine 23-Plex Assay (Bio-Rad, Hercules, CA, USA) and the Luminex system (Bio-Rad; software v6.0). All protein samples were normalised to the same concentration in sample diluent prior to assay. Cytokine concentrations are expressed as pg/mg protein. Cytokines with values above or below the standard curve range were excluded from analyses leaving 16 cytokines for analysis in placental samples. Additionally, placenta from female fetuses exhibited levels of IL-17a and G-CSF below the assay’s lower limit of detection, and therefore data presented for these cytokines only reflect values in male placentae. Samples were run in duplicate and mean %CV (range) between duplicates across all 16 cytokines was 5.7 (3.74-9.65).

### Statistics

Outcome measures were tested for normality and unequal variances (Levene test). Data that were non-normal were transformed to achieve normality, where possible. Differences between dietary groups for outcome measures were determined by ANOVA with Tukey’s post hoc, or Welch ANOVA with Games-Howell post hoc, or Kruskal-Wallis test with Steel-Dwass for non-parametric data (p<0.05). Data are presented as means α standard deviation (SD) or median and interquartile range (IQR), with 95% confidence interval diamonds in figures. Data transformed for analyses are presented as untransformed values. Repeated measures ANOVA was performed to assess the effect of maternal diet on weight gain across pregnancy, and differences between groups at each gestational age were evaluated by Welch ANOVA (p<0.05).

## Results

### Malnutrition impacts maternal metabolism

There was an overall effect of maternal diet on weight gain across pregnancy (p=0.002). HF mothers were heavier than UN throughout pregnancy (p<0.05), and maternal UN resulted in weight faltering from E9.5 to the end of pregnancy (p<0.05 vs. CON and HF). HF fed mothers did not gain more weight than CON (Figure 1). In addition to weight faltering, UN mothers had lower glucose, insulin, leptin and triglyceride concentrations and leptin:adiponectin ratio at GD18.5 compared to CON (p<0.05; Table 1). HF dams, despite not weighing more than CON, had increased leptin concentrations and leptin:bodyweight and leptin:adiponectin ratios, and reduced triglycerides and adiponectin concentrations at GD18.5 compared to CON (p<0.01; Table 1).

**Table 1.** Maternal metabolic profile at GD18.5.

|  | CON | UN | HF | p-value |
| --- | --- | --- | --- | --- |
| Glucose (mmol/l) | 9.5 ± 1.1<sup>a</sup> | 7.5 ± 0.85<sup>b</sup> | 8.5 ± 1.3<sup>ab</sup> | 0.014 |
| Insulin (ng/ml) | 0.59 ± 0.24<sup>a</sup> | 0.13 ± 0.06<sup>b</sup> | 0.41 ± 0.18<sup>ab</sup> | 0.008 |
| Leptin (ng/ml) | 145 (116-167)<sup>a</sup> | 1.0 (0.68-2.0)<sup>b</sup> | 174 (147-1175)<sup>ab</sup> | <0.0001 |
| Leptin:body weight | 4.0 (3.1-4.6)<sup>a</sup> | 0.04 (0.03-0.08)<sup>b</sup> | 5.2 (4.0-31.0)<sup>ab</sup> | <0.0001 |
| Triglycerides (mg/dl) | 121 (97.7-149)<sup>a</sup> | 57.6 (50.6-62.1)<sup>b</sup> | 70.3 (61.2-82.4)<sup>b</sup> | <0.0001 |
| HMW Adiponectin (µg/ml) | 6.7 ± 1.1<sup>a</sup> | 7.1 ± 1.2<sup>a</sup> | 4.3 ± 1.8<sup>b</sup> | 0.0091 |
| Leptin:adiponectin ratio | 24.5 (14.9-28.0)<sup>b</sup> | 0.13 (0.09-0.26)<sup>c</sup> | 50.5 (41.7-160)<sup>a</sup> | 0.0009 |

Data are mean ± SD or median (IQR). Groups with different letters are significantly different (Tukey's, Steel-Dwass or Games-Howell *post hoc*). GD = gestational day (term = 19 days). HMW = high molecular weight.

**Figure 1.**
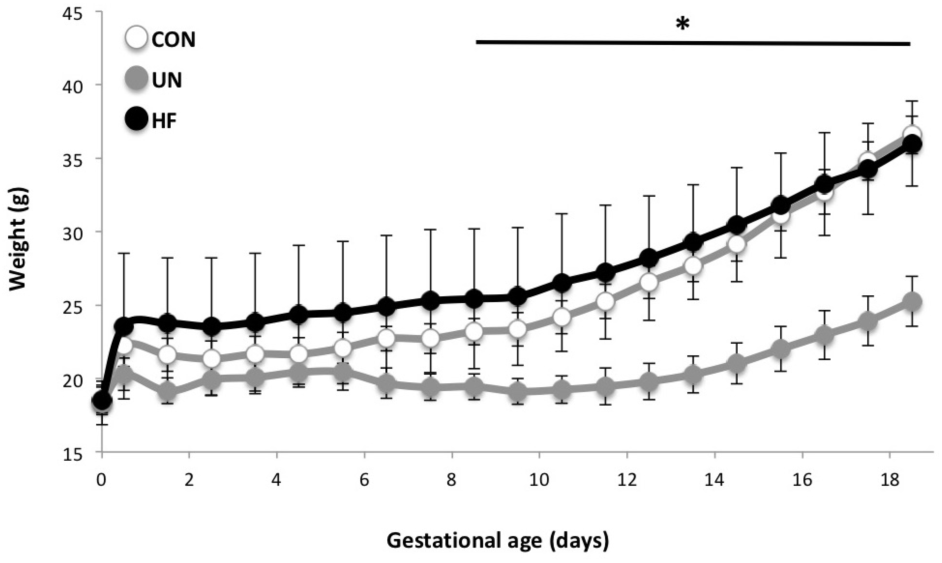
Maternal weight gain during pregnancy. There was an overall effect of maternal diet on weight gain across pregnancy (p=0.002). HF mothers were heavier than UN throughout pregnancy (p<0.05), and maternal UN resulted in weight faltering from GD9.5 to the end of pregnancy (*p<0.05 vs. CON and HF). Data are mean +/-SD.

### Poor maternal diet has little effect on placental inflammatory markers

There were no major differences in placental inflammation between dietary groups across a panel of cytokines and chemokines, although TNF-α levels were slightly elevated in UN placentae (p=0.04; Supplementary Table 2).

### Poor maternal diet impacts fetoplacental growth and is associated with altered placental architecture

Maternal diet did not alter litter size or number of fetal resorptions at GD18.5 (Table 2). However, fetal weight, placental weight and fetal weight to placental weight ratio were significantly lower in UN compared to both CON and HF pregnancies (p<0.01, Table 2). When stratifying analyses by fetal sex, patterns in fetoplacental weights remained: both male and female fetuses of UN mothers were significantly lighter than CON and HF fetuses; placental weights of UN male fetuses were significantly lighter than CON male placentae (p<0.05, Table 2).

**Table 2.** Pregnancy outcomes at GD18.5.

|  | CON | UN | HF | p-value |
| --- | --- | --- | --- | --- |
| Fetal resorptions (n) | 1 (0-1.0) | 1 (0-2.0) | 1 (1-1.8) | NS |
| Litter size (n) | 8.3 ± 0.95 | 7.6 ± 1.1 | 7.6 ± 1.1 | NS |
| Fetal weight (g) (male and female) | 1.2 (1.1-1.3)<sup>a</sup> | 0.70 (0.66-0.98)<sup>b</sup> | 1.2 (1.1-1.3)<sup>a</sup> | <0.0001 |
| Males (g) | 1.2 (1.1-1.3)<sup>a</sup> | 0.69 (0.67-0.99)<sup>b</sup> | 1.2 (1.1-1.3)<sup>a</sup> | 0.001 |
| Females (g) | 1.2 (1.1-1.3)<sup>ab</sup> | 0.74 (0.62-0.99)<sup>b</sup> | 1.2 (1.1-1.3)<sup>a</sup> | 0.03 |
| Placental weight (g) (male and female) | 0.12 ± 0.01<sup>a</sup> | 0.10 ± 0.01<sup>b</sup> | 0.12 ± 0.01<sup>a</sup> | 0.005 |
| Males (g) | 0.13 ± 0.02<sup>a</sup> | 0.10 ± 0.01<sup>b</sup> | 0.12 ± 0.01<sup>ab</sup> | 0.007 |
| Females (g) | 0.11 ± 0.02 | 0.10 ± 0.02 | 0.11 ± 0.02 | NS |
| Fetal:placental weight (male and female) | 9.9 ± 1.9<sup>a</sup> | 7.8 ± 1.1<sup>b</sup> | 10.6 ± 1.7<sup>a</sup> | 0.0007 |
| Fetal glucose (mmol/l) (male and female) | 1.7 (1.3-1.7) | 2.3 (1.3-3.8) | 1.6 (1.3-1.6) | NS |

Data are mean ± SD or median (IQR). Groups with different letters are significantly different (Tukey's, Steel-Dwass or Games-Howell *post hoc*).

Both the junctional and labyrinth areas of UN and HF placentae were smaller compared to controls; differences were greater than 10%, where UN placentae had significantly reduced JZ area compared to CON (p=0.03; Figure 2). In-depth analysis of placental architecture revealed significantly smaller calculated labyrinthine size in HF placentae compared to CON (p=0.03, Figure 3). There was no effect of maternal diet on placental JZ size or JZ size:labyrinthine size ratio. Additionally, analysis of placental architecture revealed significantly reduced fetal blood space area in HF placentae compared to CON and UN placentae (p=0.003, Table 3 and Figure 3). Sex-stratified analyses of placental architecture showed similar trends in both male and female placentae, but results were only significant in males (p=0.001, Table 3). Fetal blood space:fetal weight ratio was significantly reduced in HF fetuses compared to CON and UN (p<0.0001, Figure 4); sex-stratified analyses showed reduced fetal blood space:fetal weight ratio in both male (p=0.004) and female (p=0.001) HF placentae compared to CON (Figure 4).

**Table 3.** Placental architecture areas in CON, UN and HF placentae at GD18.5.

| Morphological structure area (µm<sup>2</sup>) | CON | UN | HF | p-value |
| --- | --- | --- | --- | --- |
| **Spongiotrophoblast** |  |  |  |  |
| Pooled (male and female) | 435 ± 63.6 | 425 ± 103 | 413 ± 30.9 | NS |
| Males | 442 ± 86.5 | 429 ± 96.3 | 393 ± 14.8 | NS |
| Females | 428 ± 42.6 | 420 ± 123 | 433 ± 31.1 | NS |
| **Labyrinthine trophoblast** |  |  |  |  |
| Pooled (male and female) | 253 ± 7.5 | 219 ± 43.4 | 245 ± 50.9 | NS |
| Males | 255 ± 8.6 | 221 ± 45.8 | 230 ± 30.0 | NS |
| Females | 285 ± 91.4 | 217 ± 47.8 | 261 ± 66.9 | NS |
| **Trophoblast giant cells** |  |  |  |  |
| Pooled (male and female) | 27.6 ± 11.7 | 34.5 ± 14.2 | 34.0 ± 11.1 | NS |
| Males | 36.5 ± 10.2 | 40.6 ± 3.7 | 35.2 ± 12.6 | NS |
| Females | 18.6 ± 1.5 | 28.3 ± 18.9 | 32.8 ± 11.1 | NS |
| **Fetal blood space** |  |  |  |  |
| Pooled (male and female) | 148 ± 34.6<sup>a</sup> | 155 ± 34.0<sup>a</sup> | 88.1 ± 16.3<sup>b</sup> | 0.0003 |
| Males | 163 ± 39.2<sup>a</sup> | 178 ± 22.1<sup>a</sup> | 82.5 ± 8.1<sup>b</sup> | 0.0013 |
| Females | 133 ± 25.7 | 132 ± 28.0 | 93.7 ± 21.7 | NS |
| **Junctional zone sinus** |  |  |  |  |
| Pooled (male and female) | 56.6 (47.7-86.4) | 37.2 (27.9-78.2) | 61.1 (36.3-94.4) | NS |
| Males | 69.3 ± 18.3 | 62.9 ± 30.8 | 62.4 ± 26.5 | NS |
| Females | 50.7 (27.9-83.4) | 29.8 (20.1-41.7) | 66.7 (31.8-97.6) | NS |
| **Maternal labyrinthine sinus** |  |  |  |  |
| Pooled (male and female) | 317 ± 51.7 | 268 ± 59.3 | 290 ± 51.1 | NS |
| Males | 342 ± 46.2 | 290 ± 66.6 | 295 ± 72.6 | NS |
| Females | 292 ± 49.4 | 246 ± 50.0 | 286 ± 27.6 | NS |
| **Junctional zone size** |  |  |  |  |
| Pooled (male and female) | 524 ± 68.5 | 506 ± 127 | 511 ± 39.7 | NS |
| Males | 548 ± 81.1 | 533 ± 114 | 491 ± 34.6 | NS |
| Females | 501 ± 53.4 | 479 ± 152 | 531 ± 37.5 | NS |
| **Labyrinthine zone size** |  |  |  |  |
| Pooled (male and female) | 735 ± 86.7<sup>a</sup> | 666 ± 42.4<sup>ab</sup> | 624 ± 87.5<sup>b</sup> | 0.028 |
| Males | 761 ± 86.5 | 689 ± 36.4 | 607 ± 101 | NS |
| Females | 710 ± 91.3 | 596 ± 85.3 | 641 ± 83.1 | NS |
| **Junctional zone size: labyrinthine zone size ratio** |  |  |  |  |
| Pooled (male and female) | 0.72 ± 0.14 | 0.80 ± 0.24 | 0.83 ± 0.12 | NS |
| Males | 0.73 ± 0.19 | 0.78 ± 0.12 | 0.83 ± 0.17 | NS |
| Females | 0.71 ± 0.09 | 0.83 ± 0.30 | 0.83 ± 0.08 | NS |

Data are mean ± SD or median (IQR). Groups with different letters are significantly different. Area occupied by spongiotrophoblast, labyrinthine trophoblast, trophoblast giant cells, fetal blood space, junctional zone sinus and maternal labyrinthine sinus were measured by PAS and H&E staining in male and female placentae. $p < 0.05$ (ANOVA). Groups with different letters are significantly different (Tukey's post hoc).

**Figure 2.**
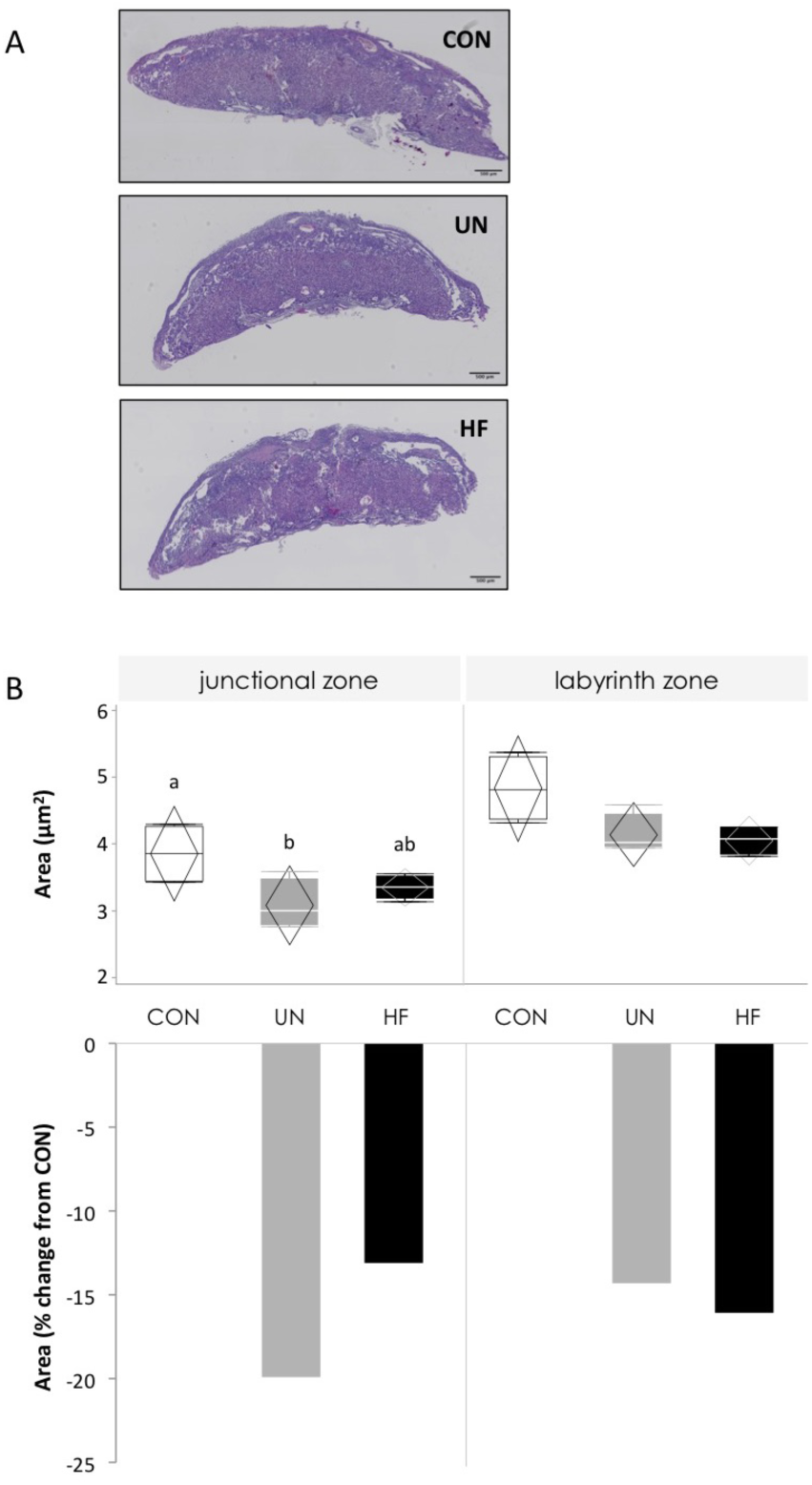
Placental morphology in CON, UN and HF placentae at GD18.5. A. Representative images of H&E stained placentae at 5X magnification. B. Area of labyrinth and junctional zones (upper panel) and percent change in area from control values (lower panel). Data are quantile box plot with 95% CI confidence diamonds.

**Figure 3.**
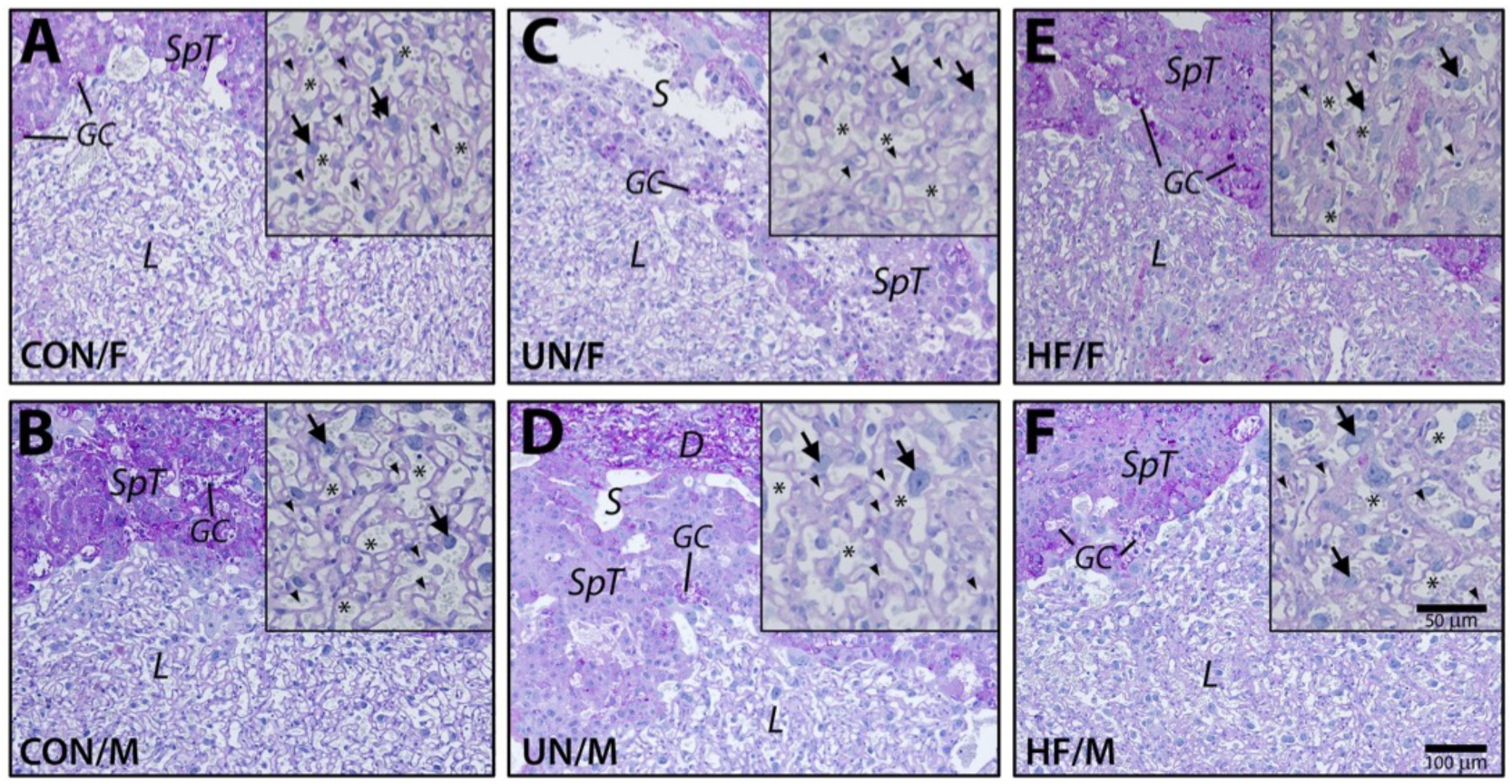
Morphology of GD18.5 placentae from male (M) and female (F) littermates of control (CON), undernourished (UN) and high fat (HF) diet pregnancies. Shown are PAS stained cross sections to detect glycogen content (red) of trophoblast glycogen cells, counterstained with hematoxylin (blue). Insets show higher magnifications of labyrinth structures. Although smaller in size and weight, UN placentae (C, D) show similar morphologies as compared to CON placentae (A, B). HF diet placentae (E, F) show more dense labyrinth structures with reduced area of fetal blood spaces. *L*, labyrinth; *SpT*, spongiotrophoblast; *GC*, glycogen cell; *S*, maternal sinusoid; *D*, decidua; arrows, sinusoidal trophoblast giant cells; arrow heads, fetal blood space; asterisk; maternal labyrinthine blood space.

**Figure 4.**
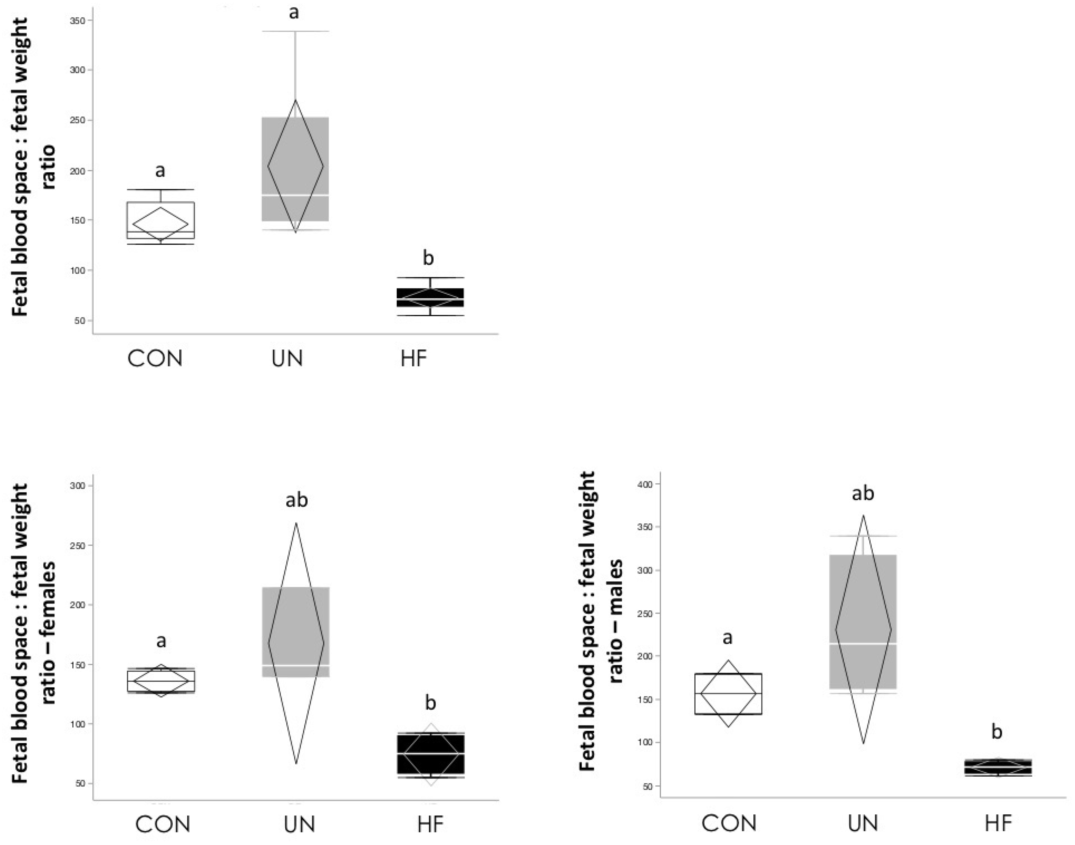
Fetal blood space:fetal weight ratio in CON, UN and HF. A. Ratio in all fetuses. B. Ratio in females. C. Ratio in males. Data are quantile box plot with 95% CI confidence diamonds. p<0.01 (Welch). Groups with different letters are significantly different (Games-Howell *post hoc*).

### Poor maternal diet impacts placental differentiation markers

We next evaluated impact of maternal diet on key placental differentiation markers (Figure 5). Secondary trophoblast giant cell marker *Ctsq*, but not *Prl3b1* (*Pl2*), mRNA expression level was lower in UN placentae compared to HF (p=0.009). mRNA expression of glycogen cell markers *Cx31.1* and *Pcdh12* were lower in HF placentae compared to UN (p=0.002) and CON (0.0003), respectively. mRNA expression of the spongiotrophoblast marker *Tpbpa* was not different between groups (Figure 5). *In situ* hybridisation for placental differentiation marker genes (*Ctsq, Prl3b1, Pcdh12* and *Tpbpa*) showed no obvious visual differences in their expression and localisation between dietary groups (Supplemental Figure 1). Ki67-positive cells were present in the chorionic plate, spongiotrophoblast and labyrinth regions of the placentae (Supplementary Figure 2). The proportion of Ki67-positive cells (to total number of cells) were significantly reduced in the chorionic plate of UN placentae compared to CON and HF (p=0.007; Supplementary Figure 2), but there were no differences between groups in other placental regions.

**Figure 5.**
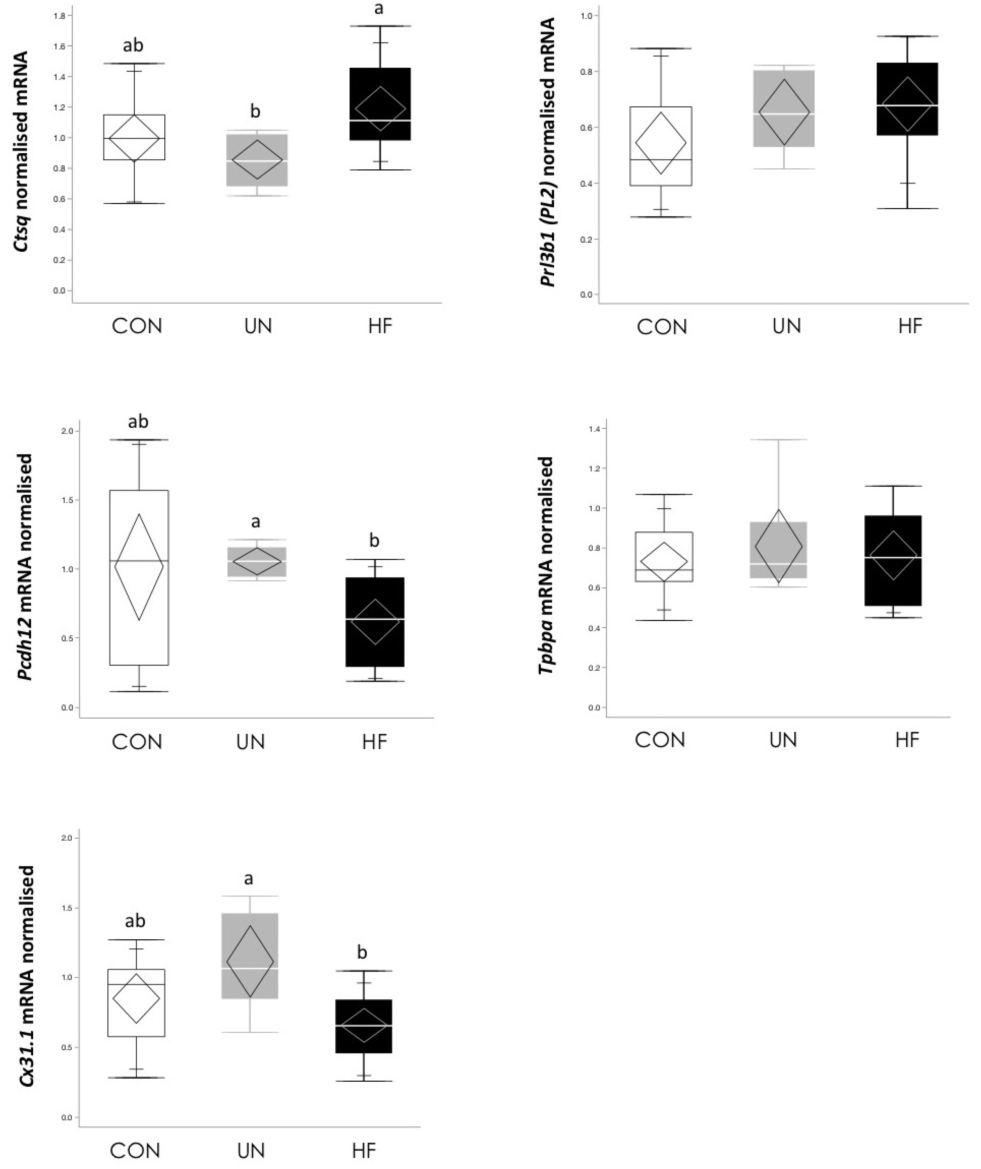
Placental differentiation marker mRNA expression levels in CON, UN, and HF male and female placentae at GD18.5. *Ctsq* mRNA expression was lower (p=0.009), and *Pcdh12* (p=0.0003) and *Cx31.1* (p=0.002) mRNA expression were higher, in UN placentae compared to HF. Data are quantile box plots with 95% CI confidence diamonds. p<0.05 (ANOVA or Welch). Groups with different letters are significantly different (Tukey’s or Games-Howell *post hoc*).

### Poor maternal diet differentially affects placental expression of ABC transporters

To determine whether placental permeability and transport was altered following maternal malnutrition, we first examined expression and localisation of placental ABC transporters. *Abcb1a*, but not *Abcb1b*, mRNA expression was lower in HF placentae compared to UN, whist *Abcg2* mRNA expression was lower in UN placentae compared to HF (Figure 6A). At the protein level, P-gp was localised to cellular membranes in both the spongiotrophoblast layer (JZ) and the labyrinth layer, with some faint and diffused staining within the cytoplasm of spongiotrophoblasts (Figure 6B). BCRP localisation and staining intensity was similar to that of P-gp, with somewhat more cytoplasmic staining in the labyrinthine layer and spongiotrophoblasts (Figure 6B). There were no differences in mean staining intensity for either protein in the spongiotrophoblast or labyrinth layers by semi-quantitative assessment (Supplementary Figure 3).

**Figure 6.**
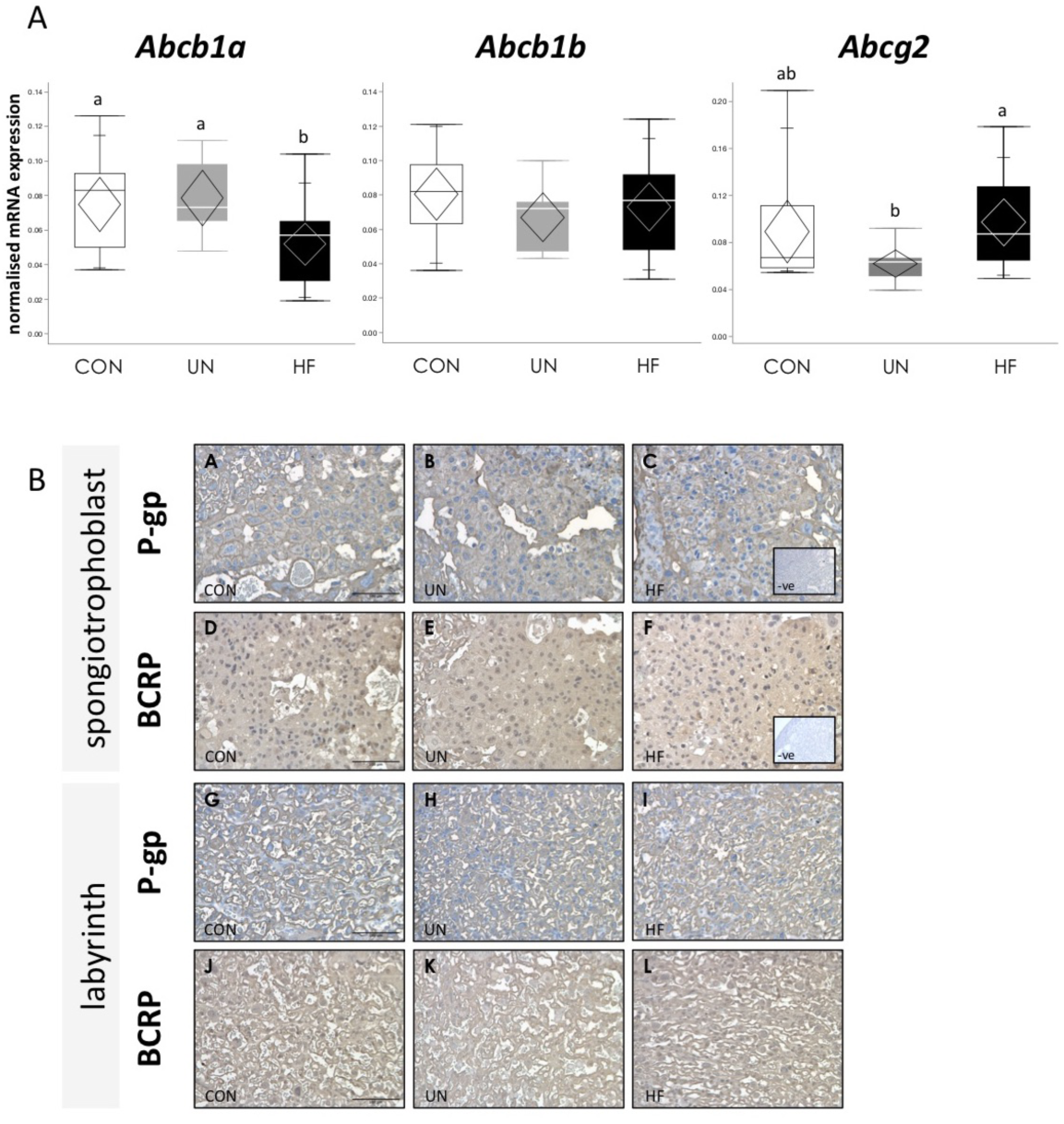
Placental ABC mRNA and protein expression in in CON, UN, and HF at GD18.5. A. Placental ABC mRNA expression levels in male and female placentae. *Abcb1a* mRNA expression was lower (p=0.012) in HF placentae compared to CON and UN and *Abcg2* mRNA expression was lower in UN placentae compared to HF (p=0.038). Data are quantile box plots with 95% CI confidence diamonds. p<0.05 (ANOVA). Groups with different letters are significantly different (Tukey’s *post hoc*). B. Representative images of immunoreactive P-gp and BCRP staining in placentae at GD18.5 in spongiotrophoblast (P-gp A-C; BCRP D-F) and labyrinth (P-gp G-I; BCRP K-L) with negative controls (inset). Images captured at 20X. Scale bar 100 µm.

### Poor maternal diet affects placental expression of fatty acid transport

There was no effect of maternal diet on mRNA expression of lipases (*El* and *Lpl*) or fatty acid transport proteins *Fatp1* and *Fatp4* (Figure 7). Fatty acid translocase *Fat/Cd36* mRNA expression was reduced in UN placentae compared to CON (p=0.001, Figure 7), whilst *Fabp*_*pm*_ was increased in UN placentae compared to CON and HF (p<0.001, Figure 7). Expression levels of FATP4 protein were lower in HF compared to UN (p=0.02, Figure 8) and consistent with mRNA findings, levels of FAT/CD36 protein were lower in UN compared to HF (p=0.045, Figure 8). There was no effect of maternal diet on protein levels of LPL, FATP1, or FAPBpm (Figure 8).

**Figure 7.**
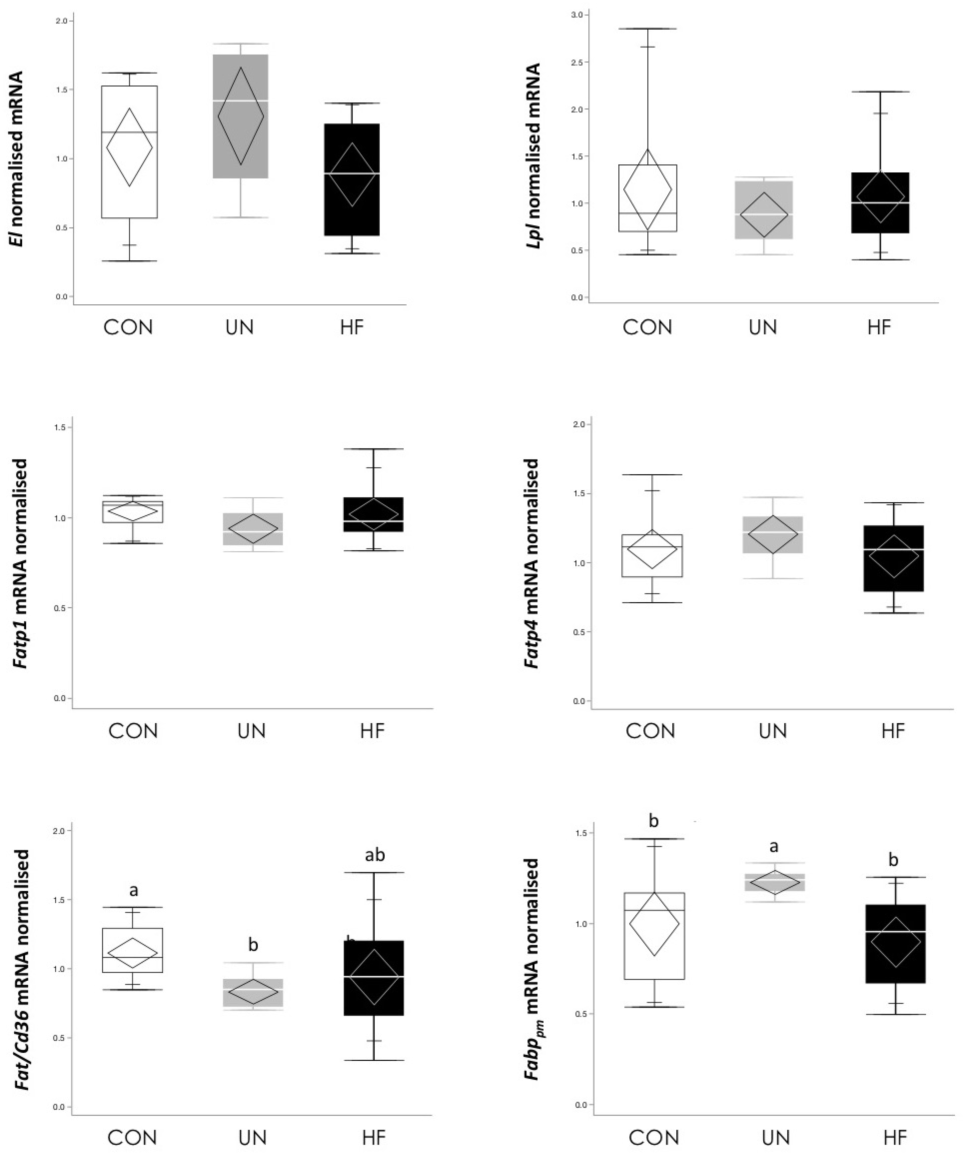
Placental fatty acid transport related marker mRNA expression levels in CON, UN, and HF male and female placentae at GD18.5. *Fat/Cd36* mRNA expression was lower in UN compared to CON placentae (p=0.001), and *Fabppm* mRNA expression was higher in UN compared to CON and HF placentae (p=0.0002.) Data are quantile box plots with 95% CI confidence diamonds. p<0.05 (ANOVA or Welch). Groups with different letters are significantly different (Tukey’s or Games-Howell *post hoc*).

**Figure 8.**
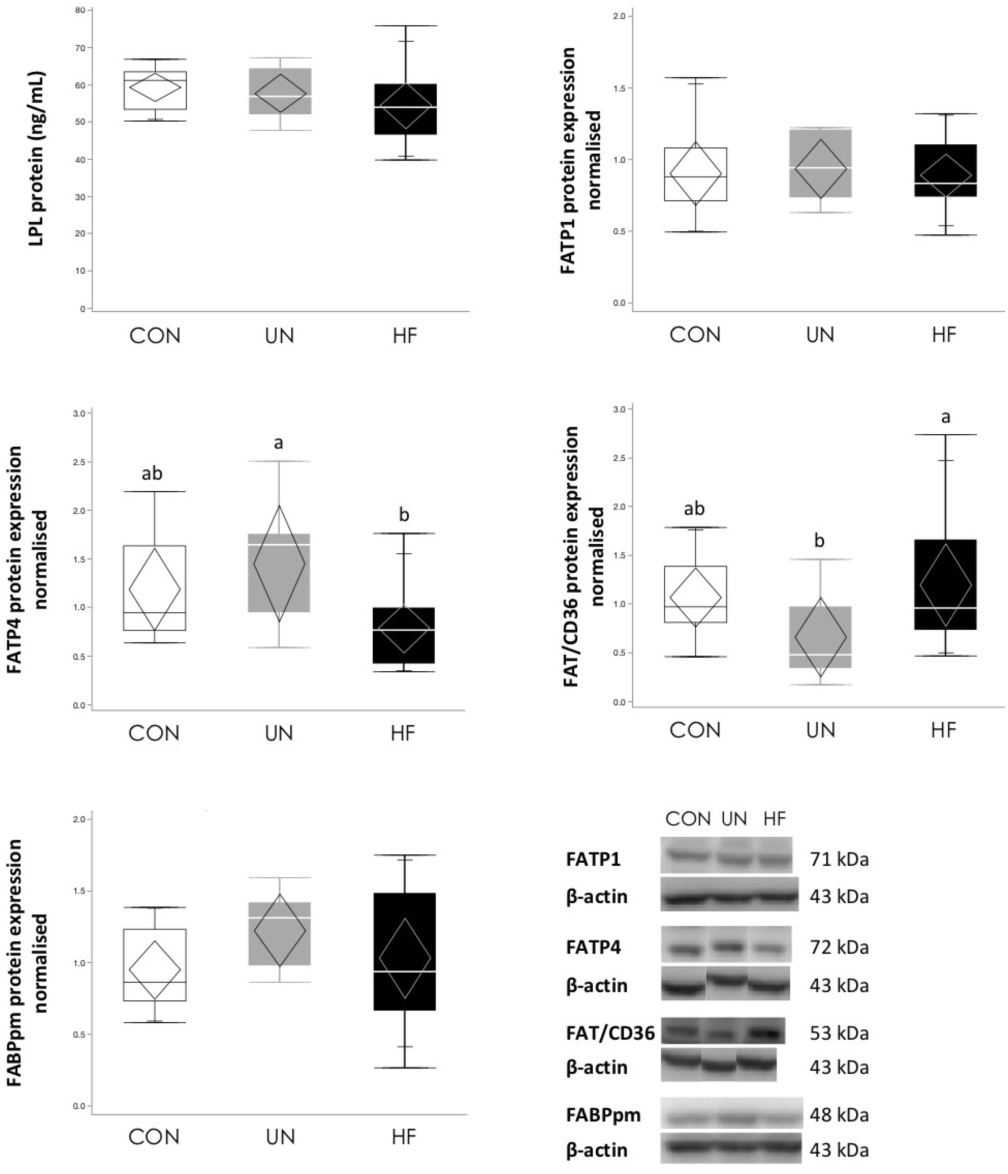
Placental fatty acid transport related marker protein expression levels in CON, UN, and HF male and female placentae at GD18.5. FAT/CD36 protein expression was lower in UN compared to HF placentae (p=0.045), and FATP4 protein expression was lower in HF compared to UN placentae (0.023). Data are quantile box plots with 95% CI confidence diamonds. p<0.05 (ANOVA). Groups with different letters are significantly different (Tukey’s *post hoc*). Representative western blots of proteins from CON, UN and HF groups in bottom right panel.

## Discussion

Appropriate placental development and function is essential to support fetal growth and development. Adequate maternal nutrition before and during pregnancy is also critical for establishing and maintaining a healthy pregnancy. Yet, it remains unclear how the placenta adapts to altered pregnancy environments, such as suboptimal maternal nutrition, in a manner that could shape fetal growth and possibly underlie early origins of later disease. We therefore undertook an approach that examined the effects of two nutritional contexts, undernutrition and overnutrition, in parallel, on placental development, and additionally sought to understand how key placental transport systems were impacted by these exposures. We hypothesised that the placenta would adapt differently to maternal UN and HF diets, through changes in its development and efflux/influx transport, and these adaptations could explain why offspring of UN and HF mothers grow differently *in utero*.

Maternal malnutrition is known to impact fetoplacental growth, the extent of which is dependent on the type and timing of the nutritional adversity^59–61^. We found that fetal and placental weights in late pregnancy were only affected by maternal UN, whilst some aspects of placental size and structure, including junctional and labyrinth zone areas, were influenced by both UN and HF diet. Overall, data from our mouse model suggest that by late pregnancy, the placenta adapted to maintain fetal growth in the face of a high fat/high calorie diet, but placental adaptations to maternal UN were insufficient to protect against fetal growth restriction. This idea is consistent with previous reports showing that maternal underweight and UN typically result in smaller fetuses/newborns, sometimes with growth restriction, and is associated with reduced placental weight and changes in placental structure^62,63^. The data are less consistent for the effects of maternal lipids^64^, increasing BMI^65–67^, and HF/high calorie diets^61,68^ on fetal growth and development: some studies report growth restriction, some report large for gestational age, and others, no effect on fetal/newborn size. We suggest that in these HF models the phenotypic outcome is largely determined by how the placenta adapts to the dietary challenge, which is in part dependent upon the timing, duration and composition of the high fat/high calorie diet relative to conception and fetal development.

To examine how the placenta may have adapted to maternal malnutrition, we took a two-pronged approach, first measuring characteristics related to its architecture and morphology. We found that UN was associated with reduced area of the JZ compared to CON. As the JZ is the major endocrine-producing layer of the mouse placenta and the TGCs help to maintain contact with the maternal environment^12^, our findings suggest that the maturation of UN JZ may be delayed, possibly impacting placental endocrine communication with the mother, including blood flow^12^. In the labyrinth, *Ctsq*, mRNA expression was reduced following UN, which may also indicate a degree of labyrinthine immaturity, with implications for exchange function. Additionally, we observed a reduction in the number of Ki67-positive cells in the chorionic plate. Reduced proliferation may be consistent with a delayed or slower growth on the fetal side of the placenta due to UN, which may compromise the flow of oxygen and nutrients to the fetus, contributing to growth restriction. Studies have found altered function, including vascular tone, in chorionic vessels of human placentae with intrauterine growth restriction^69,70^, although it is not clear if these functional deficits can be traced to earlier morphological deficiencies in the development of the chorionic plate, including changes in cellular differentiation, proliferation, or apoptosis, that have been studied only to a limited extent across gestation^71,72^.

Maternal HF diet altered the size of the labyrinthine, but not junctional zone, layers, which is in agreement with findings from another study where decidual and labyrinthine layers were reduced in placentae from HF pregnancies^73^. That study employed a somewhat different model; female mice were obese well before mating, whereas in our model, HF-fed mothers were not heavier than controls at the time of mating or throughout pregnancy. It may be that pre-pregnancy obesity has the most detrimental effect on placental architecture, which is likely to influence placental function. We also observed a significant reduction in the fetal blood space area in HF placentae, and compared to UN, reduced mRNA expression of the glycogen trophoblast cell markers *Pcdh12* and *Cx31.1.* Additionally, fetal blood space to fetal weight ratio was significantly reduced in HF placentae (in both males and females). Collectively, we believe these changes represent positive adaptations to the effects of HF diet exposure, where early and excessive growth should be avoided in the face of nutrient oversupply. As such, the HF placenta may be limiting substrate availability to the fetus by reducing the area for exchange at the level of the syncytiotrophoblast or fetal endothelial cells, the latter which may play an important, but often overlooked, roll in transplacental transport^74^. This is a hypothesis that requires further investigation. Additionally, substrate availability may be controlled through adaptations in glycogen cells, possibly due to excess nutrient supply in these HF pregnancies (and thus, reduced need to store this energy source for later use by the fetus or placenta).

Second, we measured key aspects of placental function: pro-inflammatory biomarker levels and its transport capabilities. Metainflammation, or metabolic inflammation, is thought to be a mechanism underlying developmental programming, and the placenta is a likely target for inflammatory exposures which could compromise placental development and function, and thus, impact fetal growth. Increasing maternal BMI or obesity during pregnancy has been associated with some alterations in inflammatory pathways in the mouse^73^, and human^75–77^, but not the rat^78^ placenta, but the effects of maternal underweight or undernutrition on these pathways is far less understood^3^. Our data, obtained using a comprehensive panel to measure a series of pro-inflammatory biomarkers, suggests there is no effect of maternal HF diet, and little effect of UN, on levels of pro-inflammatory molecules in the placenta near term. Our finding of increased TNF-α levels in UN placentae is consistent with increased TNF-α production^79^ and an increase in differentially expressed immune- and inflammatory-associated genes^80^ in human placentae with intrauterine growth restriction (IUGR), although we cannot definitively say that increased TNF-α in UN placentae contributes to the growth restriction seen in our fetuses. The lack of a placental inflammatory response in HF is interesting, since we have shown increased levels of pro-inflammatory cytokines in the circulation and small intestine in HF mothers^48^. Placental adaptations to maternal UN and HF diet in our model may therefore be dependent on the effects of systemic metainflammation, but independent of the effects of local (placental) inflammatory output, which may be relevant in the human context for the increasing number of pregnancies complicated by maternal overweight and increased weight gain in pregnancy, even independent of obesity^81–83^. As we did not measure inflammatory markers or gene pathways in the fetus, including organs, like the brain, known to be sensitive to inflammatory stimuli^84^, our findings do not rule out an indirect link between changes in maternal inflammatory processes with altered placental function that can drive poor fetal development. Previous studies investigating maternal obesity or HF diets on placental inflammatory pathways^73,75,76,78^ have used different models, examined fewer inflammatory markers, and/or only gene signalling pathways in the placenta, making interpretation across studies difficult. Yet, the findings from these groups and others are filling an important knowledge gap on how inflammation influences placental physiology and possibly, underlies placental pathologies. Future studies should build on the work of Kim *et al*. and Sureshchandra *et al*. to interrogate placental inflammatory pathways throughout gestation using multi-system/mulit-omics approaches in pregnancies complicated by all forms of malnourishment, not just maternal obesity.

The importance of the placenta as a conduit essential for fetal nourishment and protection is indisputable. However, most studies only investigate one aspect of its selective barrier function. Here, we investigated both, assessing select aspects of xenobiotic efflux and nutrient supply pathways. Our rationale for a more global view of transport is grounded in the reality that all forms of malnutrition often coexists with other adverse exposures like infection or inflammation, and/or medication/drug use^43–47^. Additionally, ABC transporters are sensitive to the effects of lipids^34^, but their response to undernutrition, especially in pregnancy, has been minimally investigated. Consistent with our hypothesis, transport capabilities of the UN and HF placentae were different, suggesting that at a functional level, placental adaptations may depend on the type of nutritional exposure. At the level of efflux transporters, despite that UN fetuses and placentae were smaller than CON and HF in our model, placental expression of the gene encoding the P-gp protein, *Abcb1a*, was lower in HF placentae vs. CON and UN placentae. This is inconsistent with previous findings that associated lower P-gp expression in preterm small for gestational age humans^85^ and in smaller placentae in mice^29^. However, it may be that UN placentae have adapted to provide continued protection to the UN fetus in late pregnancy, and/or these placentae remain functionally immature, given that expression levels are still high at E18.5 despite that P-gp expression has been reported to decline with advancing gestation in normal pregnancy^31,37^. That the late term UN placenta may be functionally less mature is consistent with our architectural and differentiation marker findings. Expression of the gene encoding the BCRP protein, *Abcg2*, was lower in UN vs. HF placentae, which is consistent with findings in human placenta with IUGR^86^. In that study, the authors also found when using a potent inhibitor of BCRP and at higher concentrations, other ABC proteins^87^, there was an increase in TNF-α and IFNγ-induced apoptosis in primary trophoblast and BeWo cells, which, taken with the reduced expression levels in IUGR placentae suggests that BCRP may play an important role in placental cell survival, contributing to placental dysfunction and fetal growth restriction.

The fetus, and particularly the brain, is dependent on transplacental transport of fatty acids in the later part of pregnancy to support and optimise its development, yet the exposures that influence fatty acid transfer, and how these affect fetal development, are only beginning to be understood. Here we found that malnutrition did not affect expression levels of EL and LPL, which may indicate that there is no dysfunction at the level of placental lipoprotein hydrolysis, which is required for the release of free fatty acids so that these can be taken up by the placental syncytiotrophoblasts^88,89^. However dysfunction may be occurring at the level of fatty acid transport: we found that maternal UN was associated with reduced placental FAT/CD36 mRNA and protein expression (a translocase located on both maternal and fetal sides of the trophoblast^90^) but increased *Fabp*_*pm*_ mRNA expression (located only on the maternal side). As we were unable to measure fatty acid levels in the fetus, we cannot determine at this time whether these changes in placental fatty acid transport are associated with altered circulating levels in the UN fetus or metabolism in the fetal liver, which could influence fetal growth. Still, it is likely that there is some maladaptation to maternal UN, which may prevent sufficient transfer of fatty acids to the undernourished fetus in late pregnancy, contributing to its reduced growth. In the human placenta, Chassen *et al.*^91^ showed that syncytiotrophoblast microvillous plasma membrane protein expression of FATP6 and FAT/CD36 was increased in IUGR placentae, associated with an increase in the concentrations of n-6 and n-3 long chain polyunsaturated fatty acids, which may be a result of impaired fatty acid efflux transport across the basal plasma membrane on the of the fetal-facing syncytiotrophoblast. Maternal HF diet was associated with reduced placental FATP4 protein expression (located on both maternal and fetal sides). Similar to our findings on placental architecture and development, we believe the HF placenta has appropriately adapted to the excessive nutrient supply by restraining the passage of some fatty acids in an effort to prevent fetal overgrowth, particularly excess adipose deposition^41,64^. It is important to note that our findings are limited because we investigated transporter expression in whole placental homogenates, rather than isolating the trophoblast plasma membranes. Whilst we would have had to pool multiple placentae to obtain sufficient sample for localised assessments, limiting our ability to look at individual placentae and sex differences, this is a method used in many elegant nutrient transport studies^61,92,93^, and is worth considering for future studies to truly tease apart the placental transport phenotypes associated with maternal malnutrition.

As our study is cross-sectional in design, we are unable to determine the point during which fetoplacental growth (and likewise, structure and function of the placenta) began to be impacted by these nutritional exposures, and whether the trajectory for placental development and function diverged in UN and HF pregnancies at the same time during gestation. Early placental phenotypes are important to characterise, because how the placenta responds to secondary challenges later in gestation will no doubt be influenced by how, and the rate at which, it adapts to early pregnancy exposures, and this is particularly true when speaking of its transport and selective barrier functions^41^. Future studies should examine the impact of UN and HF diets on the placental longitudinally, and relate changes in its development, structure, and function to detailed longitudinal assessments of fetal growth, body composition, and circulating lipid and inflammatory levels, when possible.

In conclusion, we have used an innovative approach to characterise and understand placental adaptations in mice to two common nutritional adversities, undernutrition and high fat/high calorie diet, in an effort to explain why fetuses grow differently depending on the environments they are exposed to in the womb. Our study fills important knowledge gaps by assessing placental architecture, development, and function, and the spectrum of malnutrition, which can impact fetoplacental development independent of maternal body composition. Additionally, our assessment of both nutrient and xenobiotic transport is critical to understand the many pregnancies globally in which multiple adversities exist, including poor nutrition, infectious disease and inflammation, and use of medications or drugs. By uncovering the relationships between adverse nutritional exposures and placental development and function, we can better understand why some fetuses are at risk for, or are protected from, compromised development. This may inform tailored nutritional approaches for women before and during pregnancy to optimise fetal development and in the longer term, postnatal growth and lifelong health.

## Acknowledgments

We thank Richard Maganga for his assistance with the animal work and Ricardo Henriques, University College London, for sharing his bioRxiv template, which we have slightly modified for use here.

## Author Contributions and Notes

Contributions: Conceptualisation, KLC, SJL, EB; methodology KLC, MK, EB; investigation, KLC, EM, MK, EB, TTNN; data curation, formal analysis, KLC, EB; writing—original draft preparation, KLC; writing— review, KLC, EB, MK, SJL, SGM, TTNN, EM. This research was funded by the Canadian Institutes of Health Research (CIHR) (grants MOP-81238 and FDN-143262 to SJL, Fellowship MFE-246638 to KLC and grant 452740 to SGM). KLC is supported by the Natural Sciences and Engineering Research Council of Canada, the Molly Towell Perinatal Research Foundation (New Investigator), Carleton University Office of Research and the CIHR. EB is funded by Conselho Nacional de Desenvolvimento Científico e Tecnológico (CNPq; 422410/2016-0).

The authors have no competing interests and nothing to disclose. This article contains supplemental figures and tables.

---

**Supplementary Figure 1.**
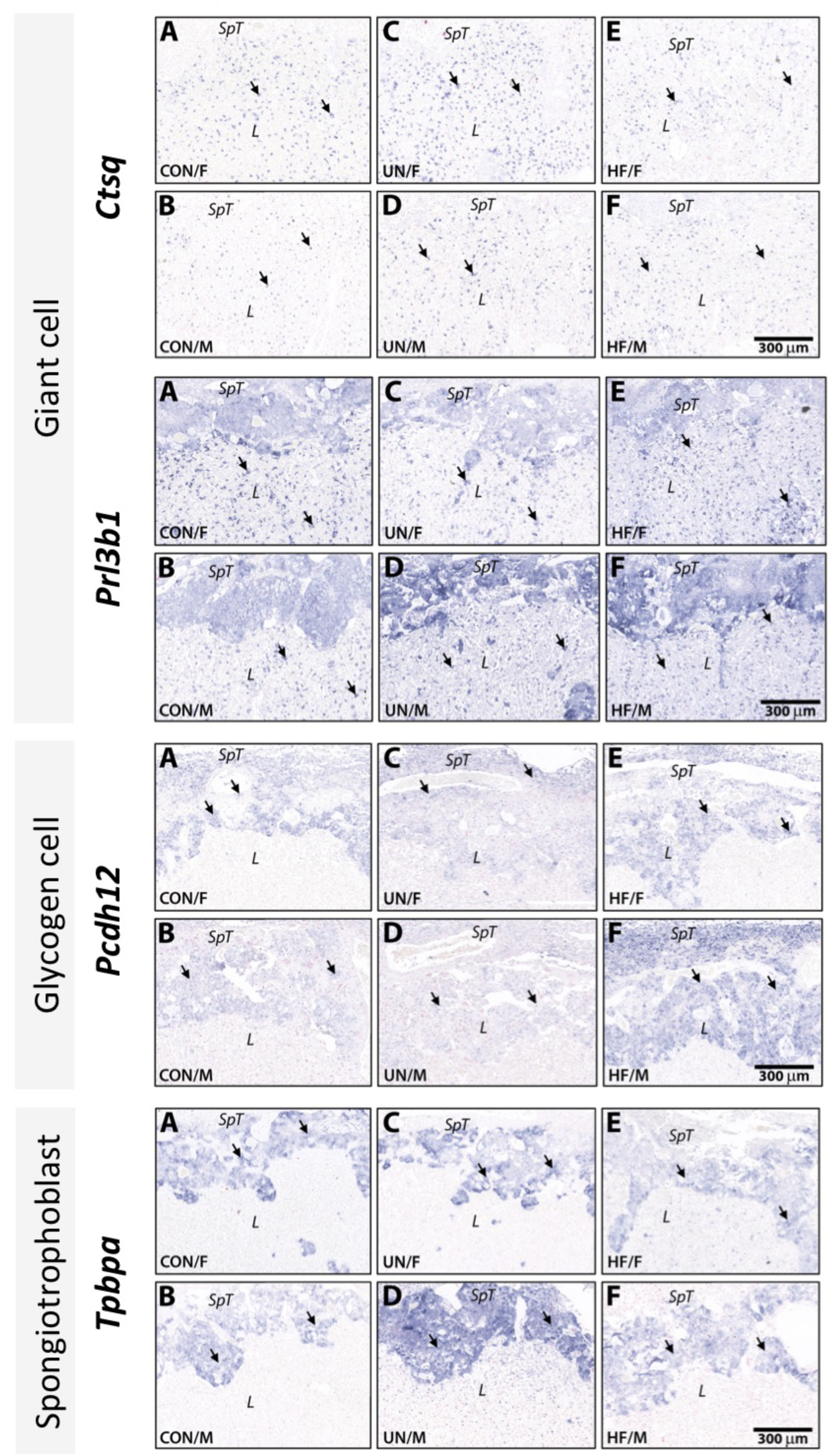
*in situ* hybridisation of GD18.5 placentae from male (M) and female (F) littermates of control (CON), undernourished (UN) and high fat (HF) diet pregnancies. Shown are cross sections probed with *Ctsq* and *Prl3b1* to detect giant cells, *Pcdh12* to detect glycogen cells, and *Tpbpa* to detect spongiotrophoblast. Although smaller in size and weight, UN placentae (C, D in all panels) show similar morphologies compared to CON placentae (A, B in all panels) and HF placentae (E, F in all panels). L, labyrinth; SpT, spongiotrophoblast; arrows, positive probe signal.

**Supplementary Figure 2.**
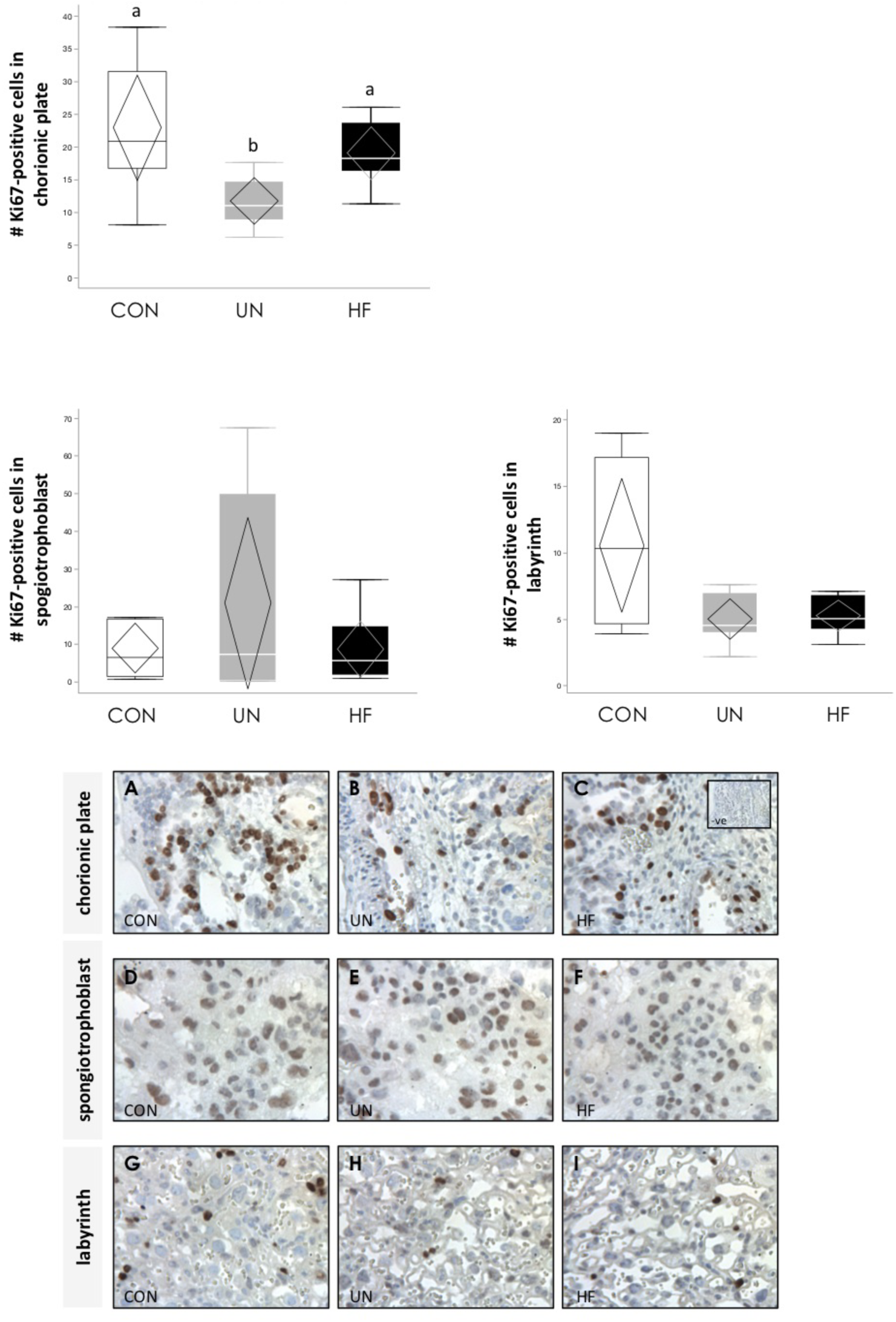
Representative images of immunoreactive Ki67-positive cell staining in CON, UN and HF placentae at GD18.5 in A. chorionic plate, B. junctional zone (spongiotrophoblast layer), and C. labyrinth zone with negative control (inset). Images captured at 40X. Data are quantile box plots with 95% CI confidence diamonds. p<0.01 (Welch). Groups with different letters are significantly different (Games-Howell *post hoc*).

**Supplementary Figure 3.**
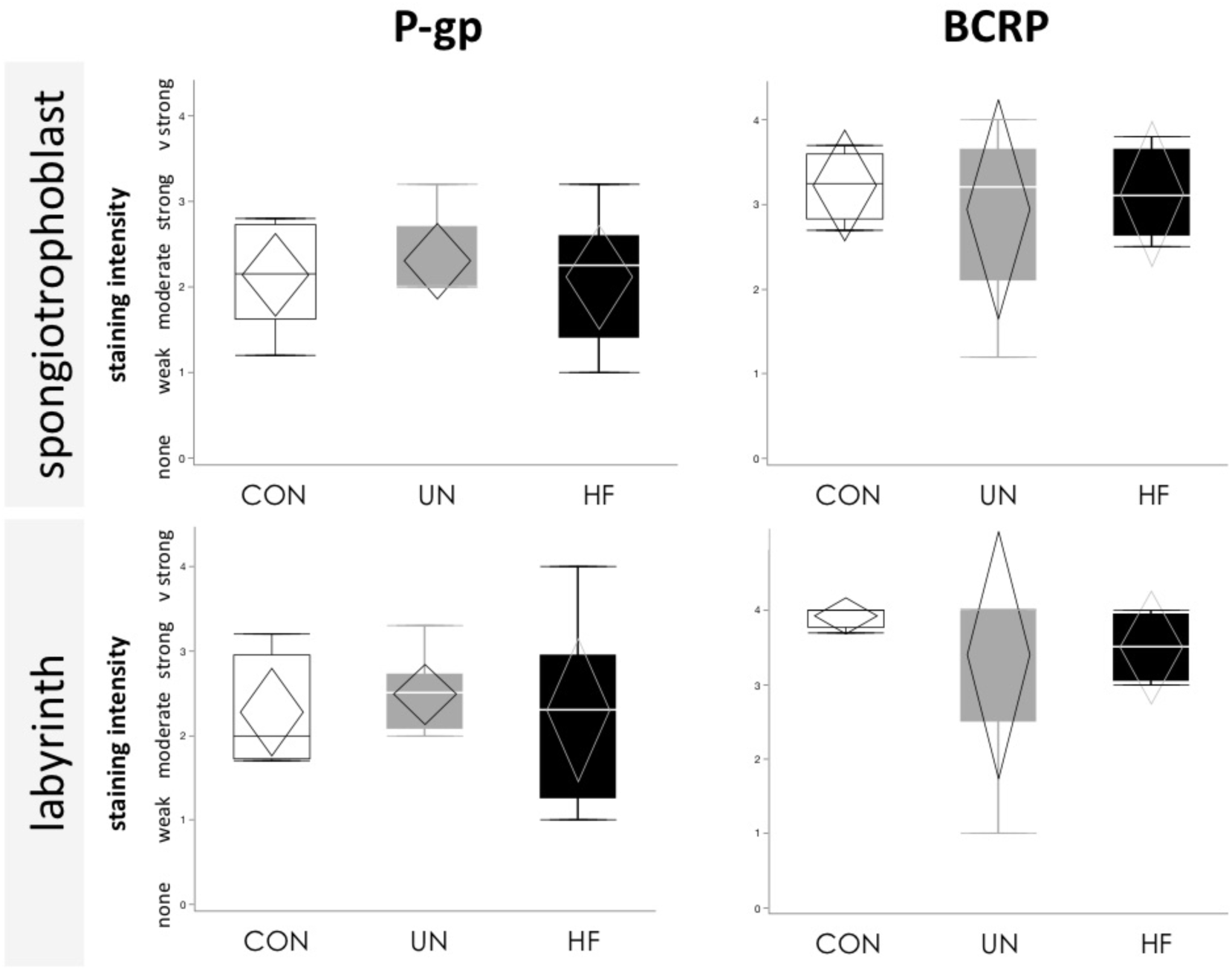
Semi-quantitative analysis of immunoreactive P-gp and BCRP staining intensity in spongiotrophoblast and labyrinth zones in CON, UN, and HF placentae at GD18.5. (refer to Figure 6B for representative images). Data are quantile box plots with 95% CI confidence diamonds.

**Supplementary Table 1.**
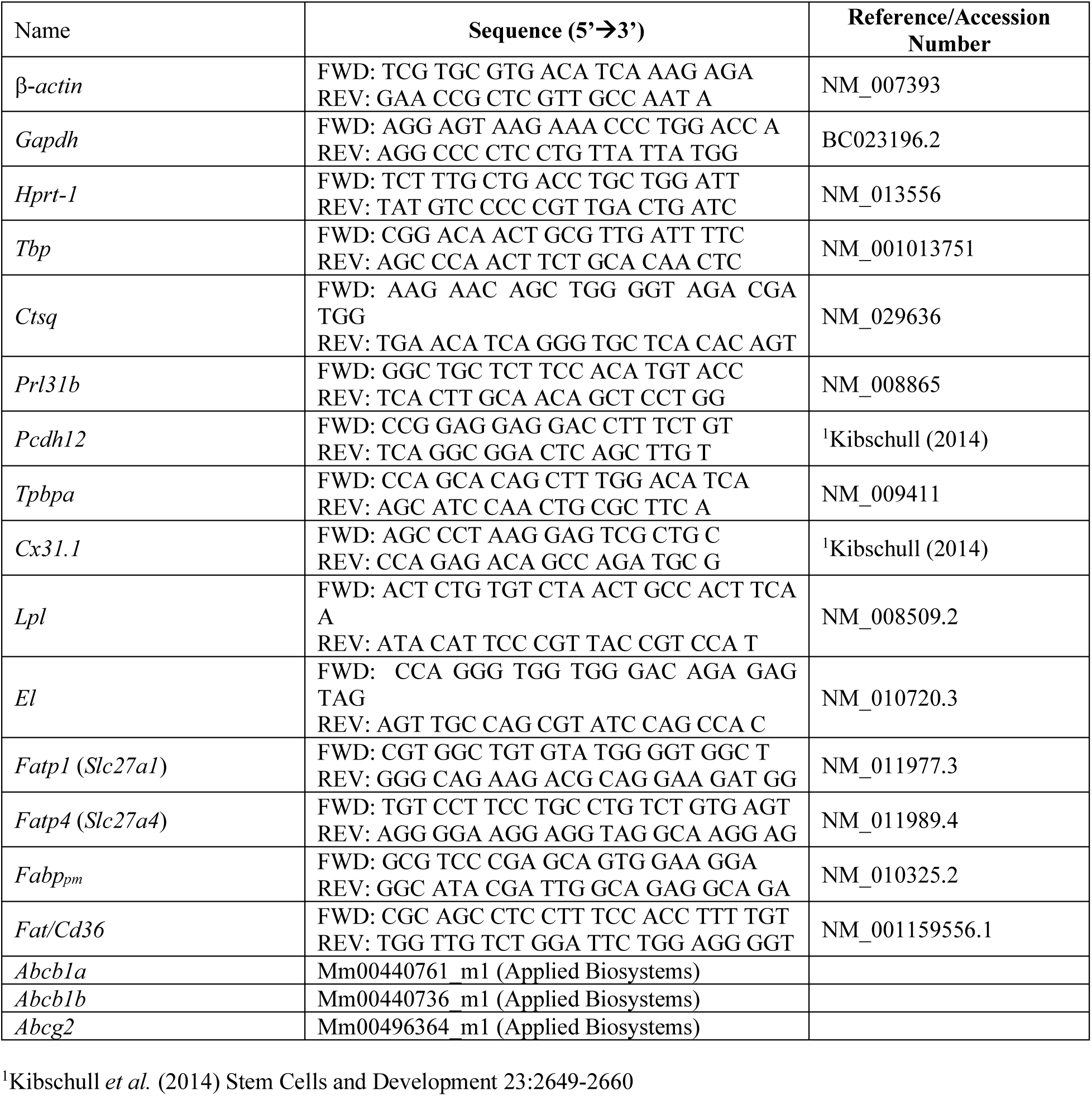
Primer sequences for qPCR.

**Supplementary Table 2.** Inflammatory cytokine and chemokine concentrations in placentae at GD18.5.

|  | CON | UN | HF | p-value |
| --- | --- | --- | --- | --- |
| IL-1α (pg/mg) | 20.9 ± 8.8 | 25.5 ± 8.4 | 22.8 ± 11.4 | NS |
| IL-1β (pg/mg) | 41.6 (38.5-46.9) | 49.9 (42.4-126) | 42.2 (39.4-43.8) | NS |
| IL-2 (pg/mg) | 2.6 ± 1.2 | 3.8 ± 1.4 | 2.7 ± 1.1 | NS |
| IL-5 (pg/mg) | 0.85 (0.74-1.0) | 1.3 (0.68-3.2) | 0.79 (0.62-0.95) | NS |
| IL-6 (pg/mg) | 0.80 (0.72-0.91) | 0.86 (0.74-2.7) | 0.84 (0.74-0.90) | NS |
| IL-10 (pg/mg) | 2.2 (2.0-3.9) | 4.3 (2.1-22.4) | 2.3 (2.0-3.8) | NS |
| IL-12p40 (pg/mg) | 1.6 (1.6-1.8) | 1.7 (1.4-4.0) | 1.6 (1.3-1.9) | NS |
| IL-12p70 (pg/mg) | 16.6 (6.6-34.8) | 24.8 (10.2-42.7) | 16.6 (9.1-36.3) | NS |
| IL-13 (pg/mg) | 5.9 (4.7-8.7) | 13.6 (8.0-44.4) | 5.4 (4.3-8.3) | NS |
| IL-17A* (pg/mg) | 0.9 (0.78-4.1) | 2.2 (1.4-4.2) | 0.97 (0.76-1.2) | NS |
| G-CSF* (pg/mg) | 2.5 (1.5-4.2) | 3.1 (2.1-5.2) | 1.6 (1.4-2.4) | NS |
| CXCL1 (KC)<br>(pg/mg) | 2.6 (1.9-8.8) | 8.9 (1.5-10.1) | 5.1 (1.4-10.5) | NS |
| MCP-1 (pg/mg) | 10.3 (6.7-15.4) | 9.8 (6.1-26.7) | 9.7 (4.2-15.1) | NS |
| MIP-1α (pg/mg) | 4.4 (3.1-5.3) | 4.0 (3.0-5.7) | 3.4 (2.3-4.1) | NS |
| MIP-1β (pg/mg) | 4.1 ± 0.69 | 3.9 ± 0.87 | 3.6 ± 0.56 | NS |
| TNF-α (pg/mg) | 31.5 (19.3-52.9)<sup>a</sup> | 50.3 (37.8-132)<sup>a</sup> | 27.8 (22.8-49.8)<sup>a</sup> | 0.044 |

Data are mean ± SD or median (IQR). Groups with different letters are significantly different (Steel-Dwass *post hoc*). \*For these inflammatory markers, only data from male placentae included because data from all female placentae were below the limit of detection of the assay. IL = interleukin; G-CSF = granulocyte-colony stimulating factor; CXCL1 (KC) = chemokine (C-X-C motif) ligand 1; MCP-1 = monocyte chemoattractant protein 1; MIP = macrophage inflammatory protein; TNF = tumor necrosis factor.

